# T-dependent B cell responses to *Plasmodium* induce antibodies that form a high-avidity multivalent complex with the circumsporozoite protein

**DOI:** 10.1101/108746

**Authors:** Camilla R. Fisher, Henry J. Sutton, Joe A. Kaczmarski, Hayley A. McNamara, Ben Clifton, Joshua Mitchell, Yeping Cai, Johanna N. Dups, Nicholas J. D'Arcy, Mandeep Singh, Aaron Chuah, Thomas S. Peat, Colin J. Jackson, Ian A. Cockburn

## Abstract

The repeat region of the *Plasmodium falciparum* circumsporozoite protein (CSP) is a major vaccine antigen because it can be targeted by parasite neutralizing antibodies; however, little is known about this interaction. We used isothermal titration calorimetry, X-ray crystallography and mutagenesis-validated modeling to analyze the binding of a murine neutralizing antibody to *Plasmodium falciparum* CSP. Strikingly, we found that the repeat region of CSP is bound by multiple antibodies. This repeating pattern allows multiple weak interactions of single F_AB_ domains to accumulate and yield a complex with a dissociation constant in the low nM range. Because the CSP protein can potentially cross-link multiple B cell receptors (BCRs) we hypothesized that the B cell response might be T cell independent. However, while there was a modest response in mice deficient in T cell help, the bulk of the response was T cell dependent. By sequencing the BCRs of CSP-repeat specific B cells in inbred mice we found that these cells underwent somatic hypermutation and affinity maturation indicative of a T-dependent response. Last, we found that the BCR repertoire of responding B cells was limited suggesting that the structural simplicity of the repeat may limit the breadth of the immune response.

**Author Summary:** Vaccines aim to protect by inducing the immune system to make molecules called antibodies that can recognize molecules on the surface of invading pathogens. In the case of malaria, our most advanced vaccine candidates aim to promote the production of antibodies that recognize the circumsporozoite protein (CSP) molecule on the surface of the invasive parasite stage called the sporozoite. In this report we use X-ray crystallography to determine the structure of CSP-binding antibodies at the atomic level. We use other techniques such as isothermal titration calorimetry and structural modeling to examine how this antibody interacts with the CSP molecule. Strikingly, we found that each CSP molecule could bind 6 antibodies. This finding has implications for the immune response and may explain why high titers of antibody are needed for protection. Moreover, because the structure of the CSP repeat is quite simple we determined that the number of different kinds of antibodies that could bind this molecule are quite small. However a high avidity interaction between those antibodies and CSP can result from a process called affinity maturation that allows the body to learn how to make improved antibodies specific for pathogen molecules. These data show that while it is challenging for the immune system to recognize and neutralize CSP, it should be possible to generate viable vaccines targeting this molecule.

## Introduction

Malaria caused by *Plasmodium falciparum* causes the deaths of around 430,000 people each year [1]. The most advanced vaccine candidate for malaria is the RTS,S/AS01 vaccine which consists of a truncated version of the sporozoite-surface circumsporozoite protein (CSP), packaged in a Hepatitis C core virus-like particle delivered in AS01 - a proprietary liposome based adjuvant [2]. Phase II and Phase III clinical trials have repeatedly demonstrated that the vaccine gives around 50% protection against clinical malaria in field settings for the first year following vaccination [3]. The bulk of protection is attributed to antibodies targeting the CSP repeat epitope included within the vaccine, with some contribution from CD4+ T cells [4]. It is still unclear why the antibody response to CSP is only partially protective. We lack structural information about how neutralizing antibodies bind to CSP and knowledge on the breadth and nature of the B cell response elicited.

Antibodies to CSP were first identified as potential mediators of protection following seminal studies that showed that immunization with irradiated sporozoites could induce sterile protection against live parasite challenge [5,6]. In the early 1980s, monoclonal antibodies (mAbs) isolated from mice immunized with sporozoites were found to be capable of blocking invasion of hepatocytes [7] and directly neutralizing parasites by precipitating the surface protein coat (a process known as the circumsporozoite reaction) [8]. These antibodies were then used to clone CSP, one of the first malaria antigens identified [8,9]. The N- and C-terminal domains of CSP from all *Plasmodium* species are separated by a repeat region, which was the target of the original mAbs [9-11]. In the 3D7 reference strain of *P. falciparum,* the CSP repeat has 38 asparagine-alanine-asparagine-proline (NANP)-repeats interspersed with 4 asparagine-valine-aspartate-proline (NVDP) repeats that are concentrated towards the N-terminus [12] though different isolates can contain slightly different numbers of repeats [13]. One of the most effective *P. falciparum* sporozoite neutralizing antibodies identified in these early studies was 2A10 which can block sporozoite infectivity *in vitro* [7] and in *in vivo* mouse models utilizing rodent *P. berghei* parasites expressing the *P. falciparum* CSP repeat region [14,15].

While CSP binding antibodies have been shown to be able to neutralize sporozoites and block infection, it has also been proposed that CSP is an immunological “decoy” that induces a suboptimal, but broad, T-independent immune response due to the CSP repeat cross-linking multiple B cell receptors (BCRs) [16]. However, it remains unknown if the repetitive regions of CSP can cross-link multiple BCRs as they are not as large as typical type-II T-independent antigens [17]. Moreover, the ability to induce a T-independent response does not preclude a T-dependent component to immunity as well: various oligomeric viral surface proteins can induce both short-lived T-independent responses and subsequent affinity matured IgG responses [18,19]. Furthermore, the very little published data on the sequences of CSP binding antibodies does not convincingly support activation of a broad B cell repertoire: a small study of five *P. falciparum* CSP mouse monoclonal antibodies (mAbs) identified some shared sequences [20]. In humans, a study that generated mAbs from three individuals who received RTS,S found that the three antibodies studied had distinct sequences though these all used similar heavy chains [21].

We therefore set out to test the hypothesis that the CSP repeat can bind multiple antibodies or BCRs and drive a T-independent immune response. To do this we undertook a comprehensive biophysical characterization of the 2A10 sporozoiteneutralizing antibody that binds to the CSP repeat. We found that this antibody binds with an avidity in the nano-molar range which was unexpected as previous studies using competition ELISAs with peptides predicted a micro-molar affinity [22,23]. Strikingly, isothermal titration calorimetry (ITC), structural analyses, and mutagenesis-validated modeling revealed that the CSP repeat can be bound by around six antibodies suggesting that the repeat may potentially crosslink multiple BCRs on the surface of a B cell. However, analysis of CSP-specific B cells revealed that CSP-specific B cells can enter Germinal Centers (GCs) and undergo affinity maturation contradicting the notion that the response to CSP is largely T-independent. Moreover, we found that the BCR repertoire of CSP-binding B cells is quite limited which may restrict the size and effectiveness of the immune response.

## Results

### Characterization of the thermodynamics of 2A10-antigen binding

We began our analysis by performing isothermal titration calorimetry (ITC) to understand the interaction between 2A10 and CSP. For ease of expression we used a recombinant CSP (rCSP) construct described previously which was slightly truncated with 27 repeats [24]. ITC experiments were run on the purified 2A10 antibody and the purified 2A10 antigen-binding fragment (F_AB_) fragment to test the thermodynamic basis of the affinity of 2A10 F_AB_ towards CSP. Experiments were also performed on the 2A10 F_AB_ fragment with the synthetic peptide antigen (NANP)_6_, which is a short segment of the antigenic NANP-repeat region of CSP (Table 1; Fig. 1). The binding free energies (ΔG) and dissociation constants (*K*_D_) were found to be -49.0 kJ/mol and 2.7 nM for the full 2A10 antibody with CSP, -40 kJ/mol and 94 nM for the 2A10 F_AB_ with CSP, and -36.4 kJ/mol and 420 nM for the 2A10 F_AB_ with the (NANP)_6_ peptide.

**Fig. 1.**
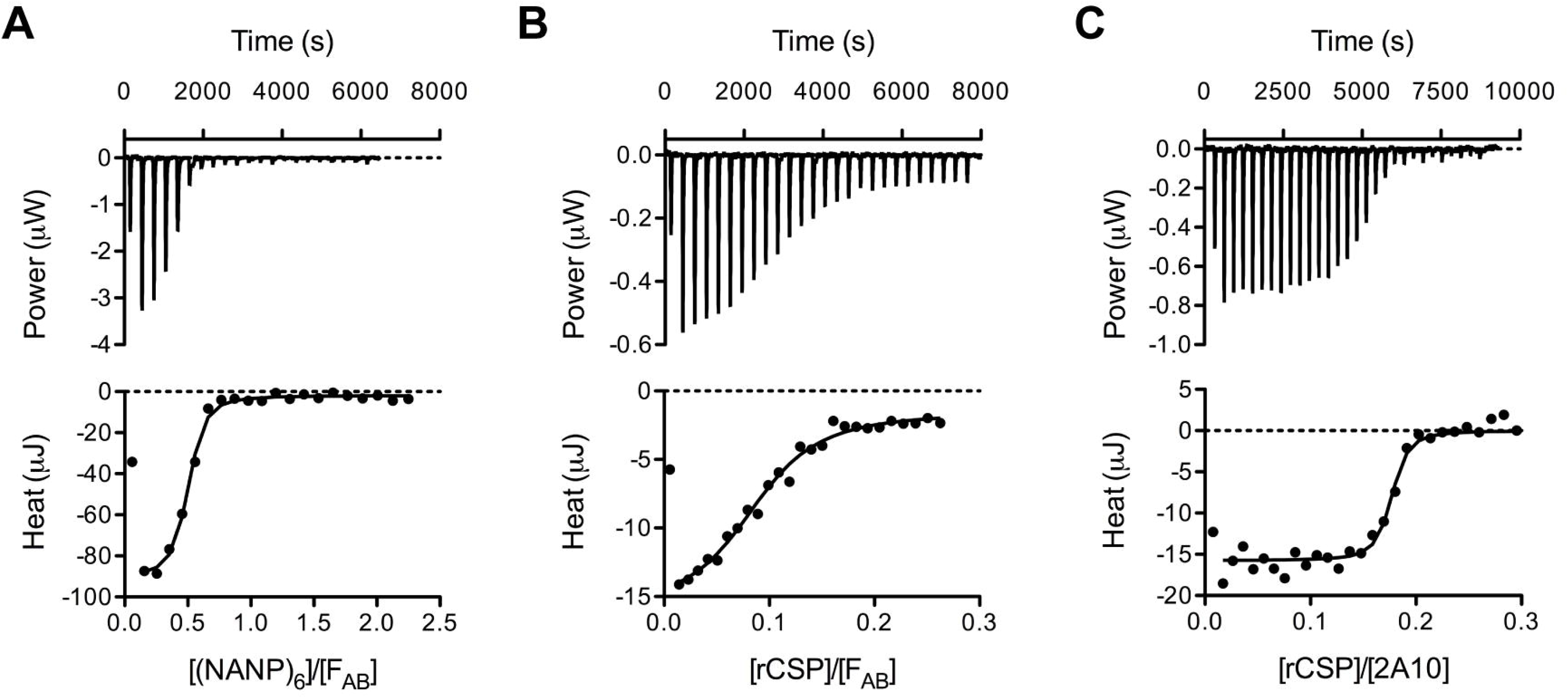
ITC data for interactions between 2A10 F_AB_ and antigens. (A) Titration of 2A10 F_AB_ with (NANP)_6_. (B) Titration of 2A10 F_AB_ with rCSP. (C) Titration of 2A10 (complete antibody) with rCSP. The upper panels represent baseline-corrected power traces. By convention, negative power corresponds to exothermic binding. The lower panels represent the integrated heat data fitted to the independent binding sites model.

**Table 1.** Thermodynamic parameters for interactions between 2A10 F_AB_, 2A10 and antigens.

| | (NANP)<sub>6</sub>:$F_{AB}$ | rCSP:$F_{AB}$ | rCSP:2A10 |
| --- | --- | --- | --- |
| $K_a$ (M<sup>-1</sup>) | $(2.37 \pm 0.91) \times 10^6$ | $(1.07 \pm 0.39) \times 10^7$ | $(3.6 \pm 2.7) \times 10^8$ |
| $K_d$ (nM) | $420 \pm 160$ | $94 \pm 34$ | $2.7 \pm 2.1$ |
| $\Delta H$ (kJ/mol complex) | $-113 \pm 5$ | $-1245 \pm 112$ | $-1175 \pm 44$ |
| $T\Delta S$ (kJ/mol complex) | $-76.6 \pm 4.9$ | $-1205 \pm 112$ | $-1126 \pm 44$ |
| $\Delta G$ (kJ/mol complex) | $-36.4 \pm 1.0$ | $-40.0 \pm 0.9$ | $-49.0 \pm 1.9$ |
| $n$ ($F_{AB}$/2A10: Ag) | $2.16 \pm 0.06$ | $10.8 \pm 0.7$ | $5.8 \pm 0.1$ |

Parameters were determined by ITC at 25 °C. Errors for $n$ (Ag : $F_{AB}$ ), $K_a$ and $\Delta H$ (complex) are 95% confidence intervals estimated from a single titration; errors for other parameters were propagated.

Surprisingly, we did not observe a typical 1:1 antibody/F_AB_ domain:antigen binding stoichiometry (Table 1). We found that each (NANP)_6_ peptide was bound to by ~2 F_AB_ fragments (2.8 repeats per F_AB_ domain). With the rCSP protein we observed that ~11 F_AB_ fragments could bind to each rCSP molecule, (2.5 repeats per F_AB_ domain. Finally, when the single-domain F_AB_ fragment is replaced by the full 2A10 antibody (which has two F_AB_ domains), we observe binding of 5.8 antibodies per rCSP molecule (4.7 repeats per antibody). Therefore all complexes exhibit approximately the same binding stoichiometry of two F_AB_ fragments/domains per ~5 repeat units. These results suggest that the antigenic region of CSP constitutes a multivalent antigen and that repeating, essentially identical, epitopes must be available for the binding of multiple F_AB_ domains.

It is not possible to separate affinity from avidity in this system, although it is apparent that there is a substantial benefit to the overall strength of binding between the antibody and antigen through the binding of multiple F_AB_ domains. The F_AB_:rCSP complex and the 2A10:rCSP complex had similar enthalpy and entropy of binding (Table 1), but the 2A10:rCSP complex had a lower overall ΔG binding, corresponding to a lower dissociation constant (2.7 nM *vs.* 94 nM for F_AB_:rCSP). The observation that this antibody-antigen (Ab-Ag) interaction is primarily enthalpically driven is consistent with the general mechanism of Ab-Ag interactions [25]. It is clear that the dissociation constant (*K*_d_) of a single F_AB_ domain to the (NANP)_6_ peptide is substantially higher (420 nM), and that the avidity, the accumulated strength of the multiple binding events between the F_AB_ domains of the antibody and the CSP repeat, is the basis for the lower *K*_d_ value observed in the 2A10:rCSP complex. Thus, the characteristic repeating pattern of the epitope on the CSP antigen allows multiple weak interactions with 2A10 F_AB_ domains to accumulate, which yields a complex with a high avidity dissociation constant in the low nM range.

### Structural analysis of the (NANP)-repeat region and the 2A10 F_AB_

To better understand the molecular basis of the multivalent interaction between 2A10 and rCSP, we performed structural analysis of the components. Previous work indicated that the NANP-repeat region of CSP adopts a flexible rod-like structure with a regular repeating helical motif that provides significant separation between the N-terminal and the C-terminal domains [26]. Here, we performed far-UV circular dichroism (CD) spectroscopy to investigate the structure of the (NANP)_6_ peptide. These results were inconsistent with a disordered random coil structure (**S1 Fig**.). Rather, the absorption maximum around 185 nm, minimum around 202 nm and shoulder between 215 and 240 nm, is characteristic of intrinsically disordered proteins that can adopt a spectrum of states [27].

The lowest energy structures of the (NANP)_6_ repeat were predicted using the PEP-FOLD *de novo* peptide structure prediction algorithm [28]. The only extended state among the lowest energy structures that was consistent with the reported spacing of the N-and C-terminal domains of CSP [26], and which presented multiple structurally similar epitopes was a linear, quasi-helical structure, which formed a regularly repeating arrangement of proline turns (Fig. 2A). The theoretical CD spectrum of this conformation was calculated (**S1 Fig.**), qualitatively matching the experimental spectra: the maximum was at 188 nm, the minimum at 203 nm and there was a broad shoulder between 215 and 240 nm. To investigate the stability of this conformation, we performed a molecular dynamics (MD) simulation on this peptide, which showed that this helical structure could unfold, and refold, on timescales of tens of nanoseconds, supporting the idea that it is a low-energy, frequently sampled, configuration in solution (**S1 Mov.**, **S2 Fig**.). We also observed the same characteristic hydrogen bonds between a carbonyl following the proline and the amide nitrogen of the alanine, and the carbonyl group of an asparagine and a backbone amide of asparagine three residues earlier, that are observed in the crystal structure of the NPNA fragment [29]. Thus, this configuration, which is consistent with previously published experimental data, is a regular, repeating, extended conformation that would allow binding of multiple F_AB_ domains to several structurally similar epitopes.

**Fig 2.**
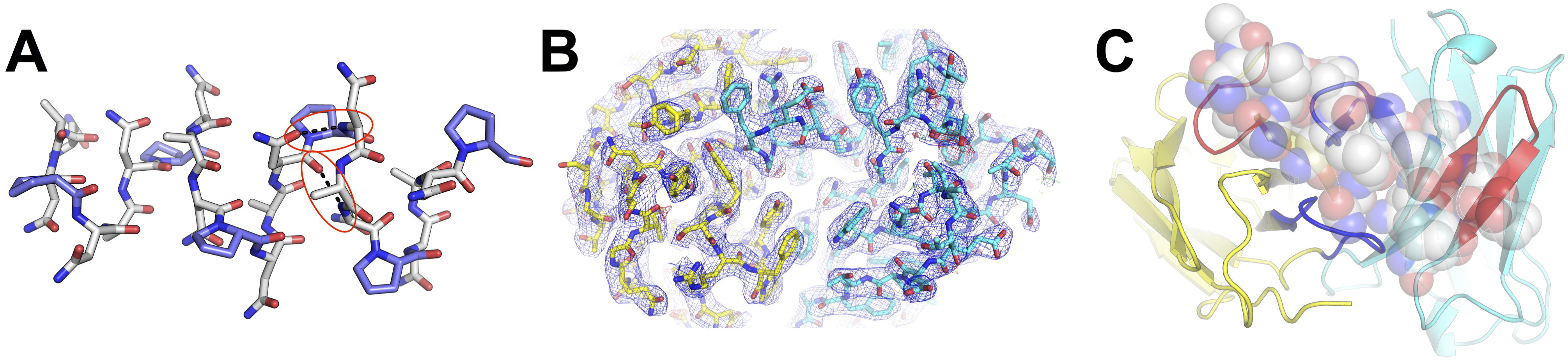
Structures of the (NANP)_6_ peptide (A), the 2A10 F_AB_ fragment (B) and the model of the F_AB_ fragment-(NANP)_6_ complex (C). (A) The calculated structure of the (NANP)_6_ peptide is a helical structure containing the same hydrogen bonds between a carbonyl following the proline and the amide nitrogen of the alanine, and the carbonyl group of an asparagine and a backbone amide of asparagine 3 residues earlier (highlighted in red) that are observed in [29]. (B) Electron density (blue mesh; 2mF_o_-dF_c_ at 1σ) of the 2A10 F_AB_ fragment viewed from above the antigen-binding site. Light chain is shown as yellow sticks, heavy chain as cyan. (C) A calculated model of the (NANP)_6_:2A10 F_AB_ fragment complex. The CDR2 regions of each chain are shown in red, the CDR3 regions of each chain are shown in blue.

To better understand the interaction between the 2A10 and the (NANP)-repeat region, we solved the crystal structure of the 2A10 F_AB_ fragment in two conditions (**S1 Table**), yielding structures that diffracted to 2.5 Å and 3.0 Å. All of the polypeptide chains were modeled in good quality electron density maps (Fig. 2B), except for residues 134-137 of the light chain. This loop is located at the opposite end of the F_AB_ fragment to the variable region and not directly relevant to antigen binding. The 2.5 Å structure contained a single polypeptide in the asymmetric unit, whereas the 3.0 Å structure contained three essentially identical chains. Superposition of the four unique F_AB_ fragments from the two structures revealed that the variable antigen binding region is structurally homogeneous, suggesting that this region might be relatively pre-organized in the 2A10 F_AB_. This is consistent with the observation that antibodies typically undergo relatively limited conformational change upon epitope binding [25]. Indeed, a recent survey of 49 Ab-Ag complexes revealed that within the binding site, the heavy chain Complementarity Determining Region (CDR)-3 was the only element that showed significant conformational change upon antigen binding and even this was only observed in one third of the antibodies [30].

Attempts to obtain a crystal structure of a complex between 2A10 F_AB_ and the (NANP)_6_ peptide were unsuccessful; unlike binary Ab-Ag interactions, in which the Ab will bind to a single epitope on an antigen and produce a population of structurally homogeneous complexes that can be crystallized, in this interaction we are dealing with an intrinsically-disordered peptide, the presence of multiple binding sites on the peptide, and the possibility that more than one 2A10 F_AB_ domain can bind the peptide. Therefore it is difficult to obtain a homogeneous population of complexes, which is a prerequisite for crystallization. Attempts to soak the (NANP)_6_ peptide into the high-solvent form of 2A10 F_AB_, in which there were no crystal packing interactions with the binding-loops, caused the crystals to dissolve, again suggesting that the heterogeneity of the peptide and the presence of multiple epitopes produces disorder that is incompatible with crystal formation.

### Modeling the interaction of the 2A10 F_AB_ with the NANP-repeat region and testing the model through site-directed mutagenesis

Although it was not possible to obtain a crystal structure of the 2A10- (NANP)_6_ peptide complex, the accurate structures of the 2A10 F_AB_ fragment, the (NANP)_6_ peptide, and the knowledge that antibodies seldom undergo significant conformational changes upon antigen binding [30], allowed us to model the interaction, which we tested using site directed mutagenesis. Computational modeling of Ab-Ag interactions has advanced considerably in recent years and several examples of complexes with close to atomic accuracy have been reported in the literature [31]. Using the SnugDock protein-protein docking algorithm [31], we obtained an initial model for binding of the peptide to the CDR region of the 2A10 F_AB_ fragment (Fig. 2C). We then performed, in triplicate, three 50 ns MD simulations on this complex to investigate whether the interaction was stable over such a time period (**S2 Mov**., **S3 Fig.**). These simulations confirmed that the binding mode that was modeled is stable, suggesting that it is a reasonable approximation of the interaction between these molecules. To experimentally verify whether our model of the 2A10 F_AB_:(NANP)_6_ peptide interaction was plausible, we performed site directed mutagenesis of residues predicted to be important for binding. Our model predicted that the interaction with (NANP)_6_ would be mainly between CDR2 and CDR3 of the light chain and CDR2 and CDR3 of the heavy chain (Fig. 2C).

In the light chain (Fig. 3A,B), Y38 is predicted to be one of the most important residues in the interaction; it contributes to the formation of a hydrophobic pocket that buries a proline residue and is within hydrogen bonding distance, *via* its hydroxyl group, to a number of backbone and side-chain groups of the peptide. Loss of this side-chain abolished binding. Y56 also forms part of the same proline-binding pocket as Y38, and loss of this side-chain also resulted in an almost complete loss of binding. R109 forms a hydrogen bond to an asparagine residue on the side of the helix; mutation of this residue to alanine in a partial loss of binding. Y116 is located at the center of the second proline-binding pocket; since loss of the entire side-chain through an alanine mutation would lead to general structural disruption of the F_AB_ fragment, we mutated this to a phenylalanine (removing the hydroxyl group), which led to a significant reduction in binding. Finally, S36A was selected as a control: the model indicated that it was outside the binding site, and the ELISA data indicated that had no effect on (NANP)_n_ binding.

**Fig 3.**
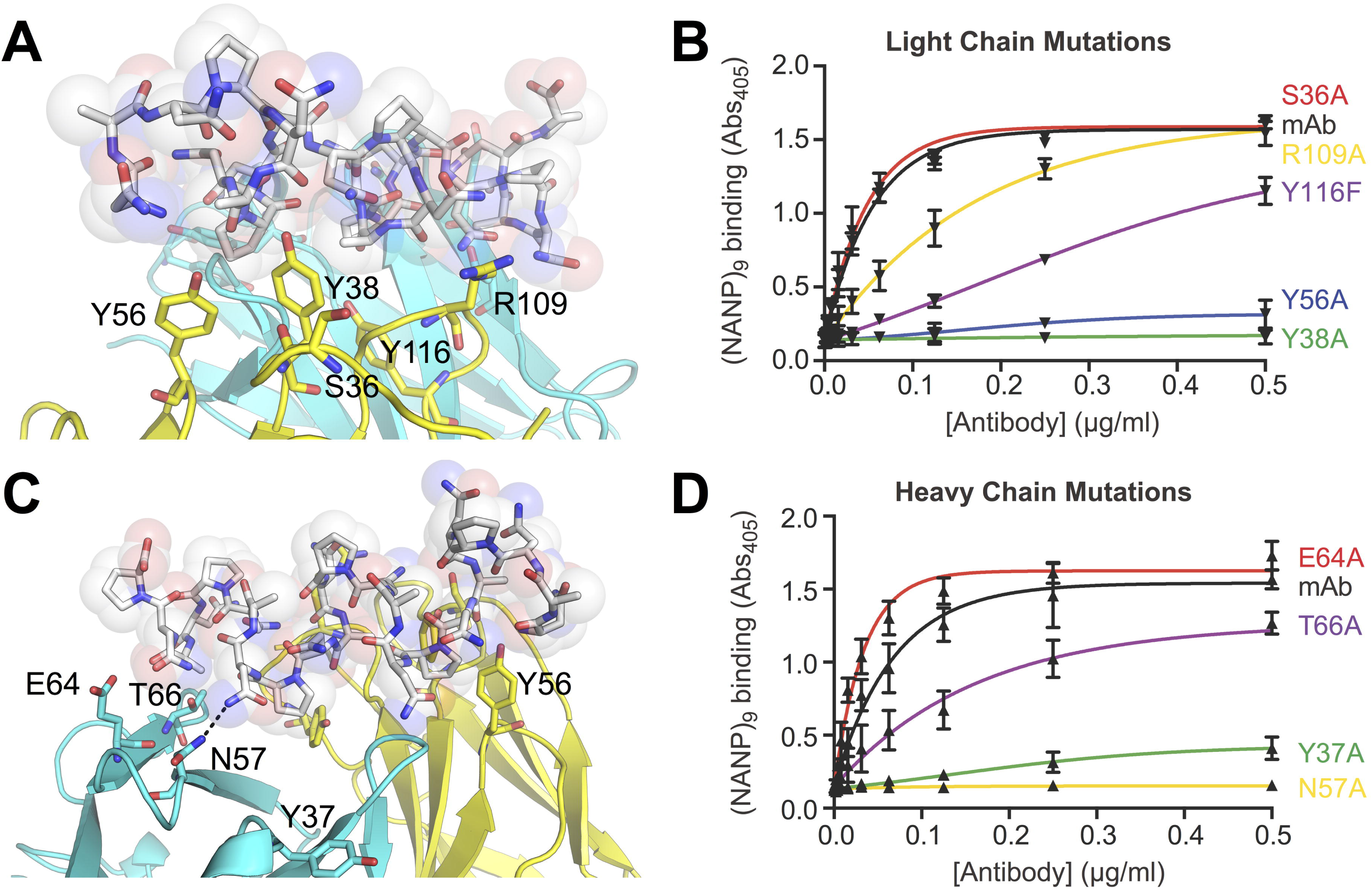
Detailed view of the (NANP)_6_:2A10 F_AB_ interface and site directed mutagenesis. (A) A model of the light chain:(NANP)_6_ interface. (B) ELISA results showing the effect of mutating light chain interface residues; error bars are based on technical replicates from one of two independent experiments. (C) A model of the heavy chain:(NANP)_6_ interface. (D) ELISA results showing the effect of mutating heavy chain interface residues; error bars are based on technical replicates from one of two independent experiments.

Within the heavy chain (Fig. 3C,D), mutation of N57 to alanine led to complete loss of binding, which is consistent with it forming a hydrogen bond to a side-chain asparagine but also being part of a relatively well packed region of the binding site that is mostly buried upon binding. T66 is located on the edge of the binding site and appears to provide hydrophobic contacts through its methyl group with the methyl side-chain of an alanine of the peptide; mutation of this residue resulted in a partial loss of binding. Interestingly, mutation of E64, which is location in an appropriate position to form some hydrogen bonds to the peptide resulted in a slight increase in binding, although charged residues on the edge of protein:protein interfaces are known to contribute primarily to specificity rather than affinity [32]. Specifically, the cost of desolvating charged residues such as glutamate is not compensated for by the hydrogen bonds that may be formed with the binding partner. Y37 is located outside the direct binding site in the apo-crystal structure; the loss of affinity could arise from long-range effects, such as destabilization of the position of nearby loops. In general, the effects of the mutations are consistent with the model of the interaction.

### The multivalency of the CSP repeat region

The binding mode of the F_AB_ fragment to the (NANP)_6_ peptide is centered on two proline residues from two non-adjacent NANP-repeats (Fig. 3A, C). These cyclic side-chains are hydrophobic in character and are buried deeply in the core of the F_AB_ antigen binding site, into hydrophobic pockets formed by Y38 and Y56 of the light chain and the interface between the two chains. In contrast, the polar asparagine residues on the sides of the helix are involved in hydrogen bonding interactions with a number of polar residues on the edge of the binding site, such as N57 of the heavy chain. Due to the twisting of the (NANP)_6_ repeat, the binding epitope of the peptide is 2.5-3 alternate NANP repeats, with a symmetrical epitope available for binding on the opposite face (Fig. 4A). Thus, this binding mode is consistent with the stoichiometry of the binding observed in the ITC measurements, where we observed a stoichiometry of two 2A10 F_AB_ fragments per (NANP)_6_ peptide. To investigate whether this binding mode was also compatible with the indication from ITC that ~10.7 2A10 F_AB_ fragments, or six antibodies (containing 12 F_AB_ domains) could bind the CSP protein (Table 1), we extended the peptide to its full length. It is notable that the slight twist in the NANP helix results in the epitope being offset along the length of the repeat region, thereby allowing binding of ten 2A10 F_AB_ fragments (Fig. 4B). Six 2A10 antibodies can bind if two antibodies interact by a single F_AB_ domain and the other four interact with both F_AB_ domains. The observation that the F_AB_ fragments bind sufficiently close to each other to form hydrogen bonds also explains the observation from the ITC that the complexes with rCSP, which allow adjacent F_AB_ fragment binding, have more favorable binding enthalpy, i.e. the additional bonds formed between adjacent F_AB_ fragments further stabilize the complex and lead to greater affinity (Table 1). Thus, the initially surprising stoichiometry that we observe through ITC appears to be quite feasible based on the structure of the NANP-repeat region of the rCSP protein and the nature of the rCSP-2A10 complex. It is also clear that the effect of antibody binding to this region would be to prevent the linker flexing between the N- and C-terminal domains and maintaining normal physiological function, explaining the neutralizing effect of the antibodies.

**Fig 4.**
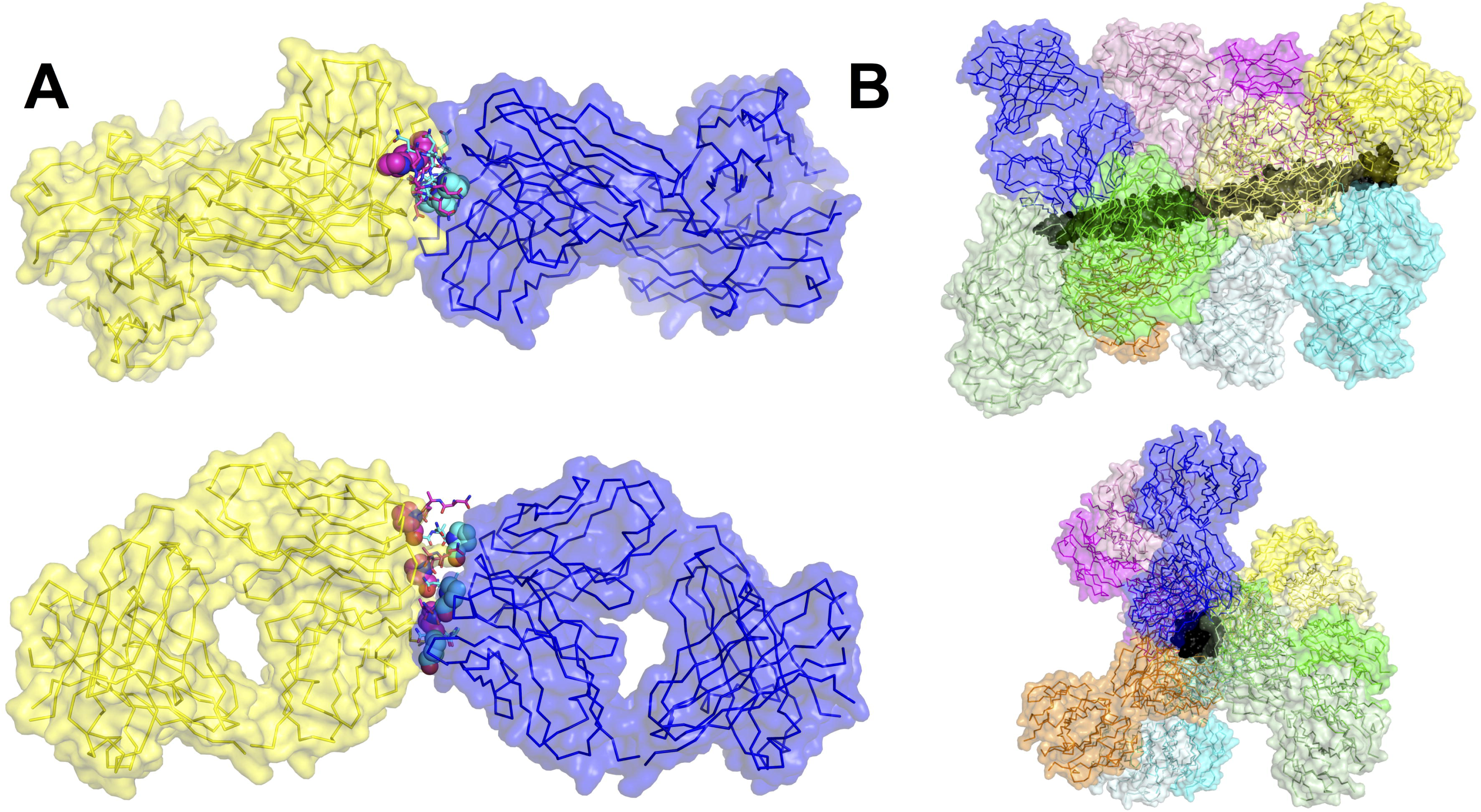
The multivalency of the NANP repeat region of the CSP protein. (A) An (NANP)_6_ peptide results in the presentation of two symmetrical epitopes, formed by alternating repeats (cyan and magenta), allowing binding by two F_AB_ domains, in keeping with the stoichiometry observed by ITC. (B) The full 27-mer repeat region results in the presentation of at least 10 separate epitopes and the twist of the helix results in displacement along the length of the repeat region, which allows binding of up to 10 separate F_AB_ fragments, consistent with 4 antibodies bound by both F_AB_ domains, and two bound by a single F_AB_ domain.

### Identification of endogenous (NANP)_n_ specific B cells to determine the BCR repertoire

We next set out to determine the implications of our structure for the B cell response to CSP. Because the CSP protein could conceivably cross-link multiple B cell receptors (BCRs) we hypothesized that the B cell response might be T-independent. As a tool to test this hypothesis we used (NANP)_n_-based tetramers to identify and phenotype antigen specific B cells in mice immunized with *P. berghei* sporozoites expressing the repeat region of the *P. falciparum* CSP (*P. berghei* CS^Pf^) [15]. The tetramers are formed by the binding of 4 biotinylated (NANP)_9_ repeats with streptavidin conjugated phycoerythrin (PE) or allophycocyanin (APC). To validate our tetramer approach, mice were immunized with either *P. berghei* CS^Pf^ or another line of *P. berghei* with a mutant CSP (*P. berghei* CS^5M^) that contains the endogenous (*P. berghei*) repeat region, which has a distinct repeat sequence (PPPPNPND)_n_. (NANP)_n_-specific cells were identified with two tetramer probes bound to different conjugates to exclude B cells that are specific for the PE or APC components of the tetramers which are numerous in mice [33]. We found that mice immunized with *P. berghei* CS^Pf^ sporozoites developed large tetramer double positive populations, which had class switched (Fig. 5A and B). In contrast, the number of tetramer double positive cells in mice receiving control parasites was the same as in unimmunized mice; moreover these cells were not class switched and appeared to be naïve precursors indicating that our tetramers are identifying bona-fide (NANP)_n_-specific cells (Fig. 5B and C). Further analysis of the different populations of B cells showed that most B cells present at this time-point were GL7^+^ CD38^-^ indicating that they are GC B cells in agreement with results from a recent publication [34] (Fig. 5B and D). Given that T cells are required to sustain GC formation beyond ~3 days these data indicate that a T-dependent response can develop to CSP following sporozoite immunization [35].

**Fig 5.**
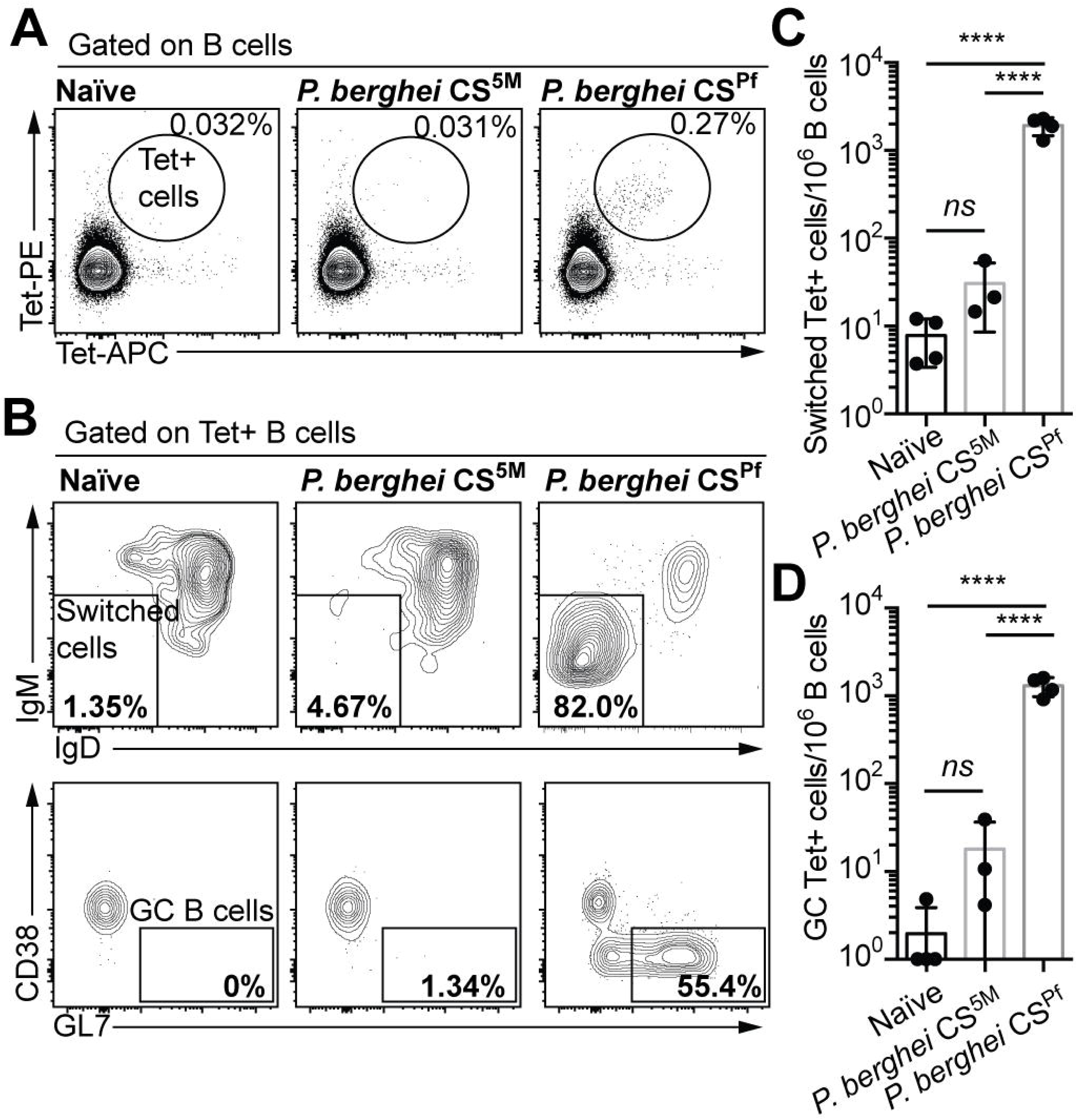
CSP-specific B cells enter the germinal center following sporozoite. BALB/C mice were immunized with either 5 × 10^4^ *P. berghei* CS^5M^ (expressing the endogenous *P. berghei* CSP repeat) or 5 × 10^4^ *P. berghei* CS^Pf^ (expressing the circumsporozoite protein from *P. falciparum*) live spoorzoites under CQ cover. 12 days later the B cell response was analyzed by flow cytometry and putative (NANP)_n_-specific cells were identified using PE and APC conjugated tetramers. (A) Representative flow cytometry plots showing the identification of (NANP)_n_-specific (Tetramer^+^) cells. (B) Representative flow cytometry plots showing the proportion of Tetramer^+^ cells that have class switched and entered a GC. (C) Quantification of the number of class switched Tetramer^+^ cells under different immunization conditions. (D) Quantification of the number of GC Tetramer^+^ cells under different immunization conditions. Data from a single representative experiment of 2 repeats, analyzed by one-way ANOVA with Tukey’s post test.

### The B cell response to the (NANP)_n_ repeat has both T-independent and T-dependent components

Our previous data showing GC formation among (NANP)_n_ specific B cells was indicative of a T-dependent response. To determine whether there might also be a T-independent component to the B cell response we immunized CD28^-/-^ mice as well as C57BL/6 controls with *P. berghei* CS^Pf^ radiation attenuated sporozoites (RAS) and measured serum (NANP)_n_ specific antibody by ELISA and the B cell response using our Tetramers. CD28^-/-^ mice have CD4+ T cells but they are unable to provide help to B cell responses [36]. Interestingly 4 days post immunization there were comparable IgM and IgG anti-(NANP)_n_ responses in the CD28^-/-^ mice and control animals (Fig. 6A), indicative of a T-independent component to immunity. However by day 27 post immunization there was no detectable IgM or IgG antibody specific for (NANP)_n_ in the CD28^-/-^ mice suggesting the T-independent response is short-lived. We further analyzed (NANP)_n_ specific B cell responses using our tetramers, in particular examining the number and phenotype (plasmablast vs GC B cell) of activated IgD^-^ Tetramer^+^ cells (Fig. 6B). In agreement with our antibody data, similar numbers of antigen specific B cells were seen at 4 days post immunization in the CD28^-/-^ and control mice and most of these cells were plasmablasts (Fig. 6C). However by 7 days post immunization the number of antigen specific cells declines in the CD28^-/-^ mice as the T dependent GC reaction begins to predominate. Thus CSP on the surface of sporozoites is able to induce short-lived T-independent B cell response, but subsequently T-dependent responses predominate.

**Fig 6.**
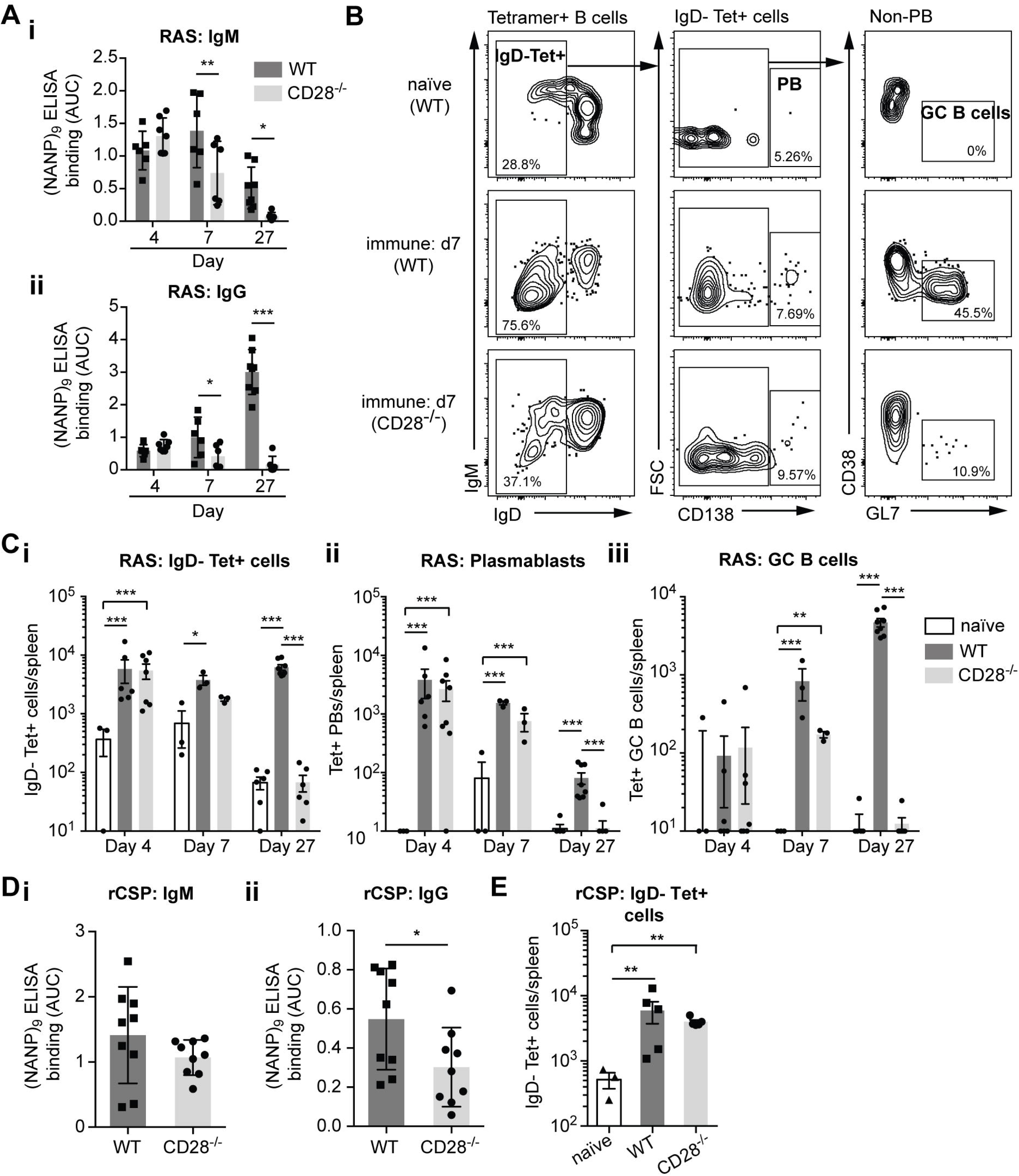
The B cell response to CSP has a T-independent component. CD28^-/-^ and control C57BL/6 mice were immunized with *P. berghei* CS^Pf^ radiation attenuated sporozoites (RAS) or rCSP in alum. Sera were taken and the spleens analyzed for antigen specific B cells using tetramers 4, 7 and 27 days post-immunization. (A) IgM and IgG (NANP)_n_ ELISA responses following RAS immunization (B) Representative flow cytometry plots 7 days post RAS immunization showing the gating of different B cell populations among Tetramer^+^ cells. (C) Absolute numbers of (i) total Tetramer^+^ IgD^-^ve (ii) Tetramer^+^ Plasmabalsts and (iii) Tetramer^+^ GC B cells post RAS immunization. (D) Antibody responses and (E) absolute numbers of Tetramer^+^ IgD^-^ B cells 4 days post immunization with rCSP. Log-transformed data pooled from 2 independent experiments for each immunization (>3 mice/group/timepoint) were analyzed using linear mixed models with day and genotype/immunization as experimental factors and the individual experiment as a random factor; only significant differences are shown.

We wanted to know if to induce a T-independent response it was necessary for CSP to be presented on the surface of the sporozoite or if free rCSP was sufficient. We found that indeed rCSP could induce a T-independent response as evidenced by similar IgM and IgG levels and IgD^-^Tetramer^+^ responses 4 days post immunization in control and CD28^-/-^ mice (Fig. 6D, E). Finally we were concerned that there may be some residual CD4^+^ T cell help in the CD28^-/-^ mice so we performed experiments in which we used the antibody GK1.5 to deplete CD4^+^ T cells [37]. In agreement with our previous data we found that sporozoites (live or RAS) and rCSP induced IgM responses in CD4 depleted mice, though we were unable to detect a significant IgG response (**Fig. S4 A**). We also detected primed antigen specific B cells in GK1.5 treated mice following RAS or rCSP immunized mice 4 days post-immunization, albeit at lower levels than in mice treated with isotype control antibodies (**Fig. S4 B**). Overall our data with GK1.5 depleted mice support our results in the CD28^-/-^ model.

### A restricted repertoire of BCRs can bind to the (NANP)_n_ repeat

Our ability to identify and sort (NANP)_n_ specific B cells with our tetramers also allows us to examine the repertoire of antibodies that can bind the (NANP)_n_ by sequencing the BCRs of the identified cells. While the repeat structure of CSP has been hypothesized to induce a broad polyclonal response [38], an alternative hypothesis is that the antigenically simple structure of the repeat epitope might only be recognized by a small number of naïve B cells. We therefore sorted (NANP)_n_^-^specific cells 35 days post immunization of BALB/C mice with sporozoites. We performed this analysis in BALB/C mice as this is the background of mice from which the 2A10 antibody was derived. We then prepared cDNA from the cells and amplified the heavy and kappa chain sequences using degenerate primers as described previously [39,40]. Heavy and light chain libraries were prepared from 4 immunized mice as well as from 3 naïve mice from which we bulk sorted B cells as controls. We obtained usable sequences from 3 of the 4 mice for both the heavy chain and kappa chain. Analysis of the heavy chain revealed that in each mouse 3 or 4 V regions dominated the immune response (Fig. 7A). The V regions identified (IGHV1-20; IGHV1-26; IGHV1-34 and IGHV5-9) were generally shared among the mice. As a formal measure of the diversity of our V region usage in the (NANP)_n_ specific cells and the bulk B cells from naïve mice we calculated the Shannon entropy for these populations. This analysis formally demonstrated that the diversity of the antigen specific B cells was significantly lower than the diversity of the repertoire in naïve mice (Fig. 7B). We further found that each V region was typically associated with the same D and J sequences even in different mice. For example, IGHV1-20 was typically associated with J4, IGHV5-9 with J4 while in different mice IGHV1-34 was variously paired with J1 or J4 (Fig. 7C). Similar results were obtained for the kappa chain with the response dominated by IGKV1-135; IGKV5-43/45; IGKV1-110; IGKV1-117 and IGKV14-111 (Fig. 7D and E). The V regions were typically paired with the same J regions even in different mice (Fig. 7F), for example IGKV5.43/45 was typically paired with IGKJ5 or IGKJ2 and IGKV1-110 was typically paired with IGKJ5, although IGKV1-135 was typically more promiscuous. One limitation of our high throughput sequencing approach is that the degenerate primers only amplified ~70% of the known IGHV and IGKV sequences in naïve mice, suggesting that we may not capture the full diversity of the response. However, comparison with the 5 published antibody sequences (**S2** and **S3 Table**) that include IGHV-1-20, IGKV5-45 and IGKV1-110 reveals that we are likely capturing the bulk of the antibody diversity. Together these data suggest that the number of B cell clones responding to CSP may be limited, potentially reducing the ability of the immune system to generate effective neutralizing antibodies.

**Fig 7.**
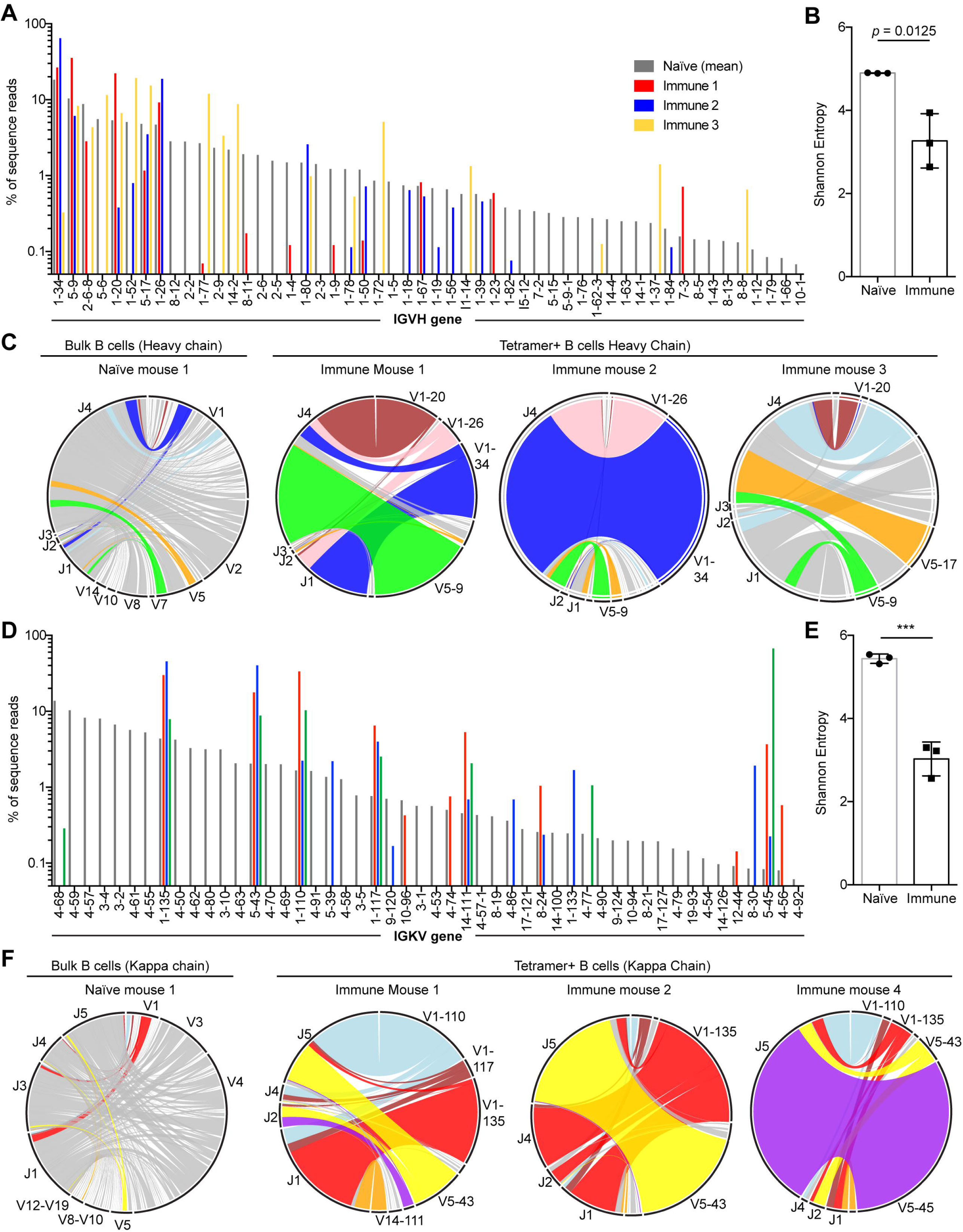
Limited diversity of (NANP)_n_ specific antibodies. BCR sequences were amplified from Tetramer^+^ cells sorted from BALB/C mice 35 days after immunization with live *P. berghei* CS^Pf^ sporozoites under CQ cover as well as bulk (B220+) B cells from naïve BALB/C mice (A) IGHV gene usage from among B cells from a representative naïve mouse (grey bars) and Tetramer+ cells from immune mice (red, blue and yellow bars). (B) Shannon’s diversity calculated for the diversity of IGHV region usage among bulk B cells and Tetramer^+^ cells. (C) Circos plots showing the IGHV-IGHJ pairings in a representative naïve mice and 3 immune mice. (D) IGKV gene usage from among B cells from a representative naïve mouse (grey bars) and Tetramer+ cells from immune mice (red, blue and green bars). (E) Shannon’s diversity calculated for the diversity of IGKV region usage among bulk B cells and Tetramer^+^ cells. (F) Circos plots showing the IGKV-IGKJ pairings in a representative naïve mouse and 3 immune mice. Statistical analysis of Shannon’s diversity index was by Student’s T test.

### CSP-binding antibodies undergo somatic hypermutation to improve affinity

Finally we were interested in knowing if the GC reaction we could see following sporozoite immunization was inducing higher affinity antibodies. We therefore examined our deep sequencing data to determine if CSP-specific antibodies had undergone somatic hypermutation (SHM) that would be indicative of B cells specific for CSP entering the GC. Taking advantage of the fact that our kappa chain primers capture the entire V-J sequences of the antibodies we sequenced we asked: 1) if the kappa chains shared between immune animals differed from the germline (providing evidence of SHM) and 2) if the mutations were conserved between different mice indicative of directed selection. Analysis of the reads from the kappa chains of the three immune mice showed that these had a much higher degree of mutation than bulk B cells from naïve mice, demonstrating SHM in the CSP-specific antibodies (Fig. 8A). We further examined each specific common kappa chain in turn (IGVK1-110; IGKV1-135; IGVK5-43/45) comparing the sequences obtained from naïve B cells and (NANP)_n_ specific cells in immune mice. This analysis showed that while, as expected, sequences from naïve mice contained few mutations, the sequences from immune mice had much higher levels of SHM. Importantly mutations were found to be concentrated in the CDR loops, and were frequently shared by immunized mice providing strong circumstantial evidence for affinity maturation (Fig. 8B; data for IGVK1-110 only shown).

**Fig 8.**
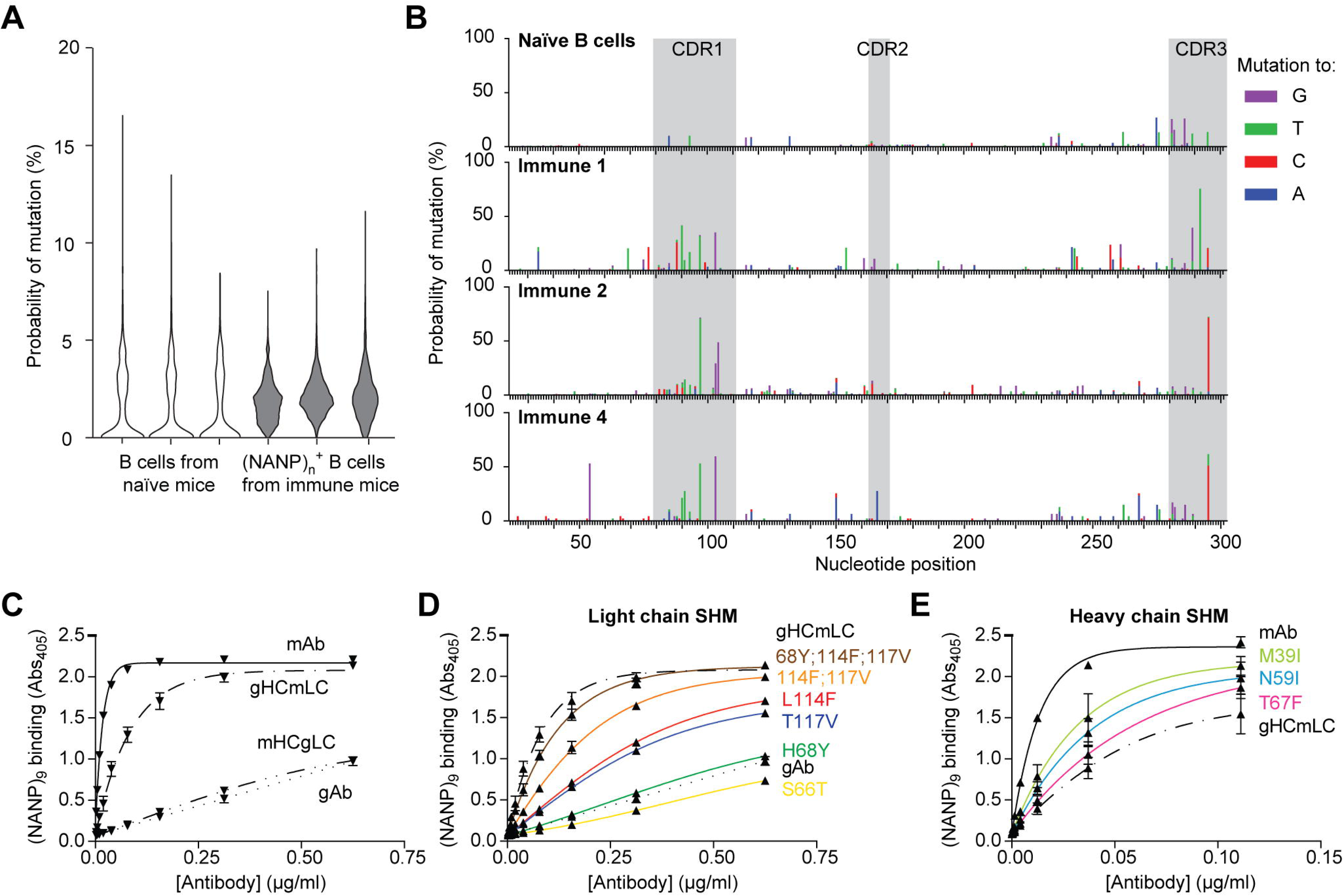
CSP-binding antibodies undergo somatic hypermutation and affinity maturation. (A) Violin plots showing the number of mutations per kappa chain read from bulk B cells from 3 individual naïve mice and sorted (NANP)_n_ specific B cells from sporozoite immunized mice (B) Skyscraper plots showing the location of mutations away from germline in the IGKV1-110 gene in a naïve mouse and in sorted (NANP)_n_ specific cells in three sporozoite immunized mice. (C) ELISA binding to the (NANP)_9_ peptide of recombinant antibodies corresponding to the 2A10 mAb, the predicted germline precursor, and hybrid antibodies containing the 2A10 heavy chain (mHC) paired with the germline light chain (gLC) and the 2010 light chain (mLC) paired with germline heavy chain (gHC). (D) Predicted mutations in the gLC were introduced to the germline precursor and their effect on binding to (NANP)_9_ measured by ELISA (E) Predicted mutations in the gHC were introduced to hybrid antibodies consisting of the mLC and the gHC heavy chain and their effect on binding to (NANP)_9_ measured by ELISA.

To directly test if CSP-binding antibodies undergo affinity maturation we expressed the predicted germline precursor to the 2A10 antibody (2A10 gAb) in HEK293T cells. We identified the predicted germline precursors of the 2A10 heavy and light chains using the program V-quest [41] (**Figs. S5** and **S6**). This analysis identified the heavy chain as IGHV9-3; IGHD1-3; IGHJ4 and the light chain as IGKV10-94;IGKJ2, with the monoclonal antibody carrying 6 mutations in the heavy chain and 7 in the light chain. The 2A10 gAb had considerably lower binding in ELISA assays compared to the 2A10 mAb itself (Fig. 8C), indicative that affinity maturation had taken place in this antibody. To determine the relative contribution of mutations in the heavy and light chain to enhancing binding we also made hybrid antibodies consisting of the mAb heavy chain and the gAb light chain and vice versa. Interestingly mutations in the light chain were almost entirely sufficient to explain the enhanced binding by the mAb compared to the gAb (Fig. 8C).

To identify the specific mutations that were important we introduced the mutations individually into the gAb light chain construct. We prioritized mutations that were shared with the 27E antibody which has previously been found to be clonally related to 2A10 having been isolated from the same mouse and which shares the same germline heavy and light chains as the 2A10 mAb [20]. We found that two mutations (L114F and T117V) in the CDR3 of the light chain appeared to account for most of the gain in binding (Fig. 8C). The effect of these antibodies appeared to be additive rather than synergistic as revealed by experiments in which we introduced these mutations simultaneously (Fig. 8D). A further mutation close to the light chain CDR2 (H68Y) also caused a modest increase in binding. As expected mutations in the heavy chains appeared generally less important for increasing binding though M39I, N59I and T67F all gave modest increases in binding (Fig. 8E). Collectively our data suggest that CSP repeat antibodies can undergo SHM in GCs resulting in affinity maturation, however the antibody response may be limited by the number of naïve B cells that can recognize and respond to this antigen.

## Discussion

Here we provide an analysis of the structure of a *Plasmodium falciparum* sporozoite-neutralizing antibody (2A10). Having obtained this structure we further modeled the binding 2A10 with its antigen target, the repeat region of CSP, and provide a thermodynamic characterization of this interaction. Finally, we used novel tetramer probes to identify and sort antigen specific B cells responding to sporozoite immunization in order to measure the diversity and maturation of the antibody response. We found that the avidity of 2A10 for the rCSP molecule was in the nanomolar range, which was much higher than the affinity previously predicted from competition ELISAs with small peptides [22,23]. This affinity is a consequence of the multivalent nature of the interaction, with up to 6 antibodies being able to bind to each rCSP molecule. Our model suggests that to spatially accommodate this binding the antibodies must surround CSP in an off-set manner, which is possible due to the slight twist in the helical structure that CSP can adopt. It is notable that the twisted, repeating arrangement of the CSP linker is the only structure that would allow binding in the stoichiometry observed through the ITC. We further found that the diversity of the antibody repertoire to the CSP repeat was limited, perhaps due to the relative simplicity of the target epitope. However, these antibodies have undergone affinity maturation to improve affinity, potentially allowing protective immune responses to develop.

Using ITC we determined the dissociation constant of 2A10 for rCSP to be 2.7 nM, which is not unusual for a mouse mAb. However it is a tighter interaction than that predicted from competition ELISAs, which predicted a micro-molar affinity [22,23]. However, these competition ELISAs were performed with short peptides rather than rCSP. Indeed, when we performed ITC with a short peptide and F_AB_ fragments we too obtained a dissociation constant in the micro-molar range (0.42 μM). The difference between the F_AB_ binding to the peptide and the tight interaction of the antibody binding to full length CSP appears to be driven by a high avidity, multivalent interaction. There is also additional enthalpic stabilization (per F_AB_ domain) in the 2A10:CSP complex, although this is partially offset by the increased entropic cost associated with combining a large number of separate molecules into a single complex. One caveat of these data is that we used a slightly truncated repeat in our recombinant CSP, however it is likely that longer repeats will have further stabilization of the interaction that could result in even higher affinity interaction between CSP and binding antibodies.

The mechanism of sporozoite neutralization remains unclear, however our structural data may provide some insights. Repeat specific antibodies can directly neutralize sporozoites (without complement or other cell mediators) in the circumsporozoite reaction [8,42]. Morevoer F_AB_ fragments alone are sufficient to block invasion [42,43]. However, it is well established that activation for complement and cell mediated immunity is important for the action of blood stage-specific antibodies [44,45]. It has also been suggested that the CSP repeat might act as a hinge allowing the N-terminal domain to mask the C-terminal domain which is believed to be important for binding to and invading hepatocytes [10]. Cleavage of this N-terminal domain is therefore required to expose the C-terminal domain and facilitate invasion [10]. Antibody binding as observed here may disrupt this process in several ways, either by opening the hinge to induce the premature exposure of the C-terminal domain. Alternatively since the repeat region is directly adjacent to the proteolytic cleavage site, anti-repeat antibodies might function by sterically hindering access of the protease to CSP, thus preventing sporozoite invasion of the hepatocyte. One possible consequence of the requirement for mutivalency to increase the avidity of the antibody, is that antibodies with different binding modes may interfere with each other limiting their effectiveness.

Our results uncovering how neutralizing antibodies bind to CSP has several implications for understanding the development of the immune response to CSP. Notably the finding that the CSP molecule can be bound by multiple antibodies/B cell receptors raises the possibility that this molecule can indeed crosslink multiple BCRs and potentially act as a type-II T independent antigen [17]. We find that indeed there is a T-independent component to the response to CSP, though T cells are required to sustain the immune response beyond day 7. As such the response to CSP appears follow a similar process to that seen for several oligomeric viral entry proteins, which induce a mix of T-independent and T-dependent responses [18,19]. It maybe that T-independent responses are driven by the density of CSP molecules on the sporozoite surface; however, rCSP can also induce a small T-independent response. This suggests that the CSP protein alone is sufficient to crosslink multiple BCRs on the B cell surface which is consistent with our structural model. Interestingly, the RTS,S/AS01 vaccine based on that contains 18 CSP repeats and does appear to induce high titers of anti-CSP antibodies which initially decline rapidly and are then more stable [4,46]. This may be consistent with the induction of a short-lived a type-II T-independent plasmablast response (accounting for the initial burst of antibodies), followed by a T-dependent response (which may be the basis of the more sustained antibody titers). The relative contributions of short-lived antibody production and long-term B cell memory to protection is an area for future investigation.

The finding of a limited repertoire in the BCR sequences specific for the (NANP)_n_ repeat contradicts previous suggestions that the response to CSP might be broad and polyclonal [38]. One explanation for this limited antibody diversity is that the antigenic simplicity of the CSP repeat region limits the range of antibodies that are capable of responding. A prior example of this is the antibodies to the Rhesus (Rh) D antigen. The RhD antigen differs from RhC by only 35-36 amino acids, resulting in the creation of a minimal B cell epitope [47]. The repertoire of antibodies that can bind this epitope are accordingly limited and mainly based on the VH3-33 gene family [48]. Another potential explanation for a limited antibody repertoire could be that the (NANP)_n_ repeat shares structural similarity with a self-antigen as is speculated to happen with meningococcus type B antigens [49], however it is not clear what this self-antigen might be. One potential outcome of this finding is that if each B cell clone has a finite burst size this may limit the magnitude of the overall B cell response.

One area for future investigation is to determine the binding modes and sporozoite neutralizing capacities of other antibodies in the response. It is clear that not all CSP-repeat binding antibodies have the same capacity for sporozoite neutralization [7]. As such the finding of a limited repertoire of responding B cells may lead to the possibility that some people have holes in their antibody repertoires limiting their ability to make neutralizing antibodies. This may explain why, while there is a broad correlation between ELISA tires of antibodies to the CSP repeat and protection following RTS,S vaccination, there is no clear threshold for protection [4].

While our work has been performed with mouse antibodies, there are major similarities between mouse and human antibody loop structure [50]. The main difference between the two species is the considerably more diverse heavy chain CDR3 regions that are found in human antibodies [51]. Consequently, this leads to a much larger number of unique clones found in humans compared to mice. However, the number of different V, D and J genes and the recombination that follows are relatively similar between humans and mice [52]. From our data it can be observed that while the BCR repertoire was restricted in the V gene usage, these different V gene populations were represented in multiple unique clones, suggesting that increasing the number of clones is unlikely to substantially increase V-region usage. Our analysis was also performed on inbred mice which may also limit repertoire diversity, however studies on the human IGHV locus reveal that in any given individual ~80% V region genes are identical between the maternal and paternal allele i.e. heterozygosity is not a major driver of human V region diversity [53,54]. It is notable that all 4 human monoclonal antibodies described to date from different volunteers share the use of the IGHV3-30 gene family [21,22], suggesting that in humans as well as mice there may indeed be a constrained repertoire of responding B cells.

Overall our data provide important insights into how the antibody response to CSP develops. Our results also help explain why relatively large amounts of antibodies are required for sporozoite neutralization and suggest that the ability to generate an effective B cell response may be limited by the very simplicity of the repeat epitope. These data support previous suggestions that CSP may be a suboptimal target for vaccination. However, we also find that CSP binding antibodies can undergo somatic hypermutation and reach high affinities. This suggests if we can develop vaccination strategies to diversify the repertoire of responding B cells and favor the GC response it may be possible to generate long-term protective immunity targeting this major vaccine candidate antigen.

## Methods

### Ethics statement

All animal procedures were approved by the Animal Experimentation Ethics Committee of the Australian National University (Protocol numbers: A2013/12; A2014/62 and A2015/76). All research involving animals was conducted in accordance with the National Health and Medical Research Council's (NHMRC) Australian Code for the Care and Use of Animals for Scientific Purposes and the Australian Capital Territory Animal Welfare Act 1992.

### Mice, Immunizations and Cell Depletions

BALB/C, C57BL/6 or CD28^-/-^ [55] mice (bred in-house at the Australian National University) were immunized IV with 5 x 10^4^ *P. berghei* CS^5M^ sporozoites expressing mCherry [56] or 5 x 10^4^ *P. berghei* CS^Pf^ sporozoites dissected by hand from the salivary glands of *Anopheles stephensi* mosquitoes. Mice were either infected with live sporozoites and then treated with 0.6mg choloroquine IP daily for 10 days or immunized with irradiated sporozoites (15kRad). For immunization with rCSP, 30ug rCSP was emulsified in Imject™ Alum according to the manufacturer’s instructions (ThermoFisher Scientific) and delivered intra-peritoneally. To deplete CD4+ T cells mice were treated with two doses of 100ug GK1.5 antibody on the 2 days prior to immunization (BioXCell); control mice received an irrelevant isotype control antibody (LTF2; BioXCell).

### Flow Cytometry and sorting

Single cell preparations of lymphocytes were isolated from the spleen of immunized mice and were stained for flow cytometry or sorting by standard procedures. Cells were stained with lineage markers (anti-CD3, clone 17A2; anti-GR1, clone RB6-8C5 and anti-NKp46, clone 29A1.4) antibodies to B220 (clone RA3-6B2), IgM (clone II/41), IgD (clone 11-26c2a), GL7 (clone GL7), CD38 (clone 90), CD138 (clone 281- 2) and (NANP)_9_ tetramers conjugated to PE or APC. Antibodies were purchased from Biolegend while tetramers were prepared in house by mixing biotinylated (NANP)_9_ peptide with streptavidin conjugated PE or APC (Invitrogen) in a 4:1 molar ratio. Flow-cytometric data was collected on a BD Fortessa flow cytometer (Becton Dickinson) and analyzed using FlowJo software (FlowJo). Where necessary cells were sorted on a BD FACs Aria I or II machine.

### Sequencing of (NANP)_n_ specific cells and BCR analysis

Single cell suspensions from the spleens of immunized mice were stained with (NANP)_n_ tetramers and antibodies to B cell markers as described in the supplementary experimental procedures. Antigen specific cells were sorted on a FACS ARIA I or II instrument prior to RNA extraction with the Arturus Picopure RNA isolation kit (Invitrogen) and cDNA preparation using the iScript cDNA synthesis kit (BioRad). BCR sequences were amplified using previously described heavy and kappa chain primers including adaptor sequences allowing subsequent indexing using the Nextera indexing kit (Illumina). Analysis was performed in house using R-scripts and the program MiXCR as described in supplementary experimental procedures.

### Binding of antibody variants

Variants of the 2A10 antibody were expressed in HEK293 T cells (a kind gift of Carola Vinuesa, Australian National University) as described in the supplemental experimental procedures. Binding to the CSP repeat was tested by ELISA and ITC using standard techniques as described in the supplementary experimental procedures.

### Statistical Analysis

Statistical analysis was performed using Prism6 (GraphPad) for simple T tests and one-way ANOVAs from single experiments. Where data were pooled from multiple experiments, analysis was performed using linear mixed models in R version 3.3.3 (R foundation for Statistical Computing). Linear mixed models are a regression analysis model containing both fixed and random effects: fixed effects being the variable/treatment under examination, whilst random effects are unintended factors that may influence the variable being measured. If significance was found from running a linear mixed model, pair-wise comparisons of the least significant differences of means (LSD) was undertaken to determine at which level interactions were occurring. Statistical significance was assumed if the *p*-value was < 0.05 for a tested difference. (ns = not significant, *= *p* < 0.5, ** = *p* < 0.01, *** = *p* < 0.001, **** = *p* < 0.0001).

### Data Deposition

Sequence data generated in this paper is deposited at the NCBI BioProject database accession number PRJNA352758. Atomic coordinates and related experimental data for structural analyses are deposited in the Protein Data Bank (PDB) with PDB codes 5ZSF and 5T0Y.

## Acknowledgments

We thank the C3 Crystallisation Centre at CSIRO for help with crystal formation and the Australian Synchrotron and beamline scientists for help with data collection. We thank Michael Devoy, Harpreet Vohra and Catherine Gillespie of the Imaging and Cytometry Facility at the John Curtin School of Medical Research for assistance with flow cytometry and multi-photon microscopy.

## Supporting Information Legends

**Supplementary Methods and Tables:** Contains extended methods; S1 Table (Data collection and refinement statistics for the crystal structures of 2A10 F_AB_ presented in this work); S2 Table (Heavy chain CDR sequences of CSP binding antibodies); S3 Table (Light chain CDR sequences of CSP binding immunoglobulins) and supplementary references.

**S1 Fig: Theoretical (A) and experimental (B) CD spectra of the (NANP)_6_ peptide.** The computational prediction of the spectra (A) was performed using DichroCalc [57], the experimental spectra was measured at 222 nm at 25 °C. A peak at 185 nm, minimum at 205 nm and shoulder between 215 and 240 nm are consistent with an intrinsically disordered, but not random coil, structure.

**S2 Fig: Cluster analysis for MD simulations of (NANP)_6_ peptide.** Conformations were clustered by concatenating the trajectory and performing a Jarvis-Patrick analysis. The clusters are sorted by their RMSD from the first cluster (starting geometry). As shown, Run 2 is stable in the starting geometry for several ns, while Run 3 diverged, then reconverged to the starting geometry, where it was stable for several ns. These data suggest the quasi-helical structure observed from the ab initio calculations is stable, and can be spontaneously sampled, on a timescale of several ns.

**S3 Fig: Cluster analysis for MD simulations of (NANP)_6_ peptide.** Molecular dynamics simulation of the (NANP)_6_:F_AB_ complex. Root mean square deviation (RMSD) of the (NANP)_6_:F_AB_ complex as a function of time. Independent simulations are shown in green, black and red.

**S4 Fig: The B cell response to CSP has a T-independent component.** Mice either treated with an anti-CD4 depleting antibody or an isotype control were immunizaed with either *P. berghei* CS^Pf^ RAS, live *P. berghei* CS^Pf^ under CQ cover or rCSP. (A) 4 days later the IgM and IgG response to the (NANP)_n_ repeat was analyzed by ELISA (B) At the same time the number of IgD^-^ Tetramer^+^ B cells was quantified in the spleen. Data are from a single experiment, analyzed using linear models with immunization/treatment as the experimental factor.

**S5 Fig: Alignment of 2A10 heavy chain and the predicted germline sequence** Residues that are mutated away from the predicted germline sequence in more one or more other antibody heavy chain (2E7 or 3D6) are highlighted in red, mutations that are predicted to be involved in binding to CSP are highlighted in blue.

**S6 Fig: Alignment of 2A10 heavy chain and the predicted germline sequence** Residues that are mutated away from the predicted germline sequence in both 2A10 and the related 2E7 antibody are highlighted in red, mutations that are predicted to be involved in binding to CSP are highlighted in blue.

**Movie S1: Molecular Dynamics simulation of the solution structure of the (NANP)_6_ peptide**

Excerpt from (NANP)_6_ run 3. The trajectory was fitted to minimize alpha-carbon RMSD and then passed through a low-pass filter with a filter length of 8 frames to reduce temporal aliasing.

**Movie S2: Molecular Dynamics simulation of the interaction of the (NANP)_n_ repeat with the 2A10 F_AB_**

Excerpt from 2A10:(NANP)_6_ run 3. The trajectory was fitted to minimize alpha-carbon RMSD and then passed through a low-pass filter with a filter length 8 frames to reduce temporal aliasing.

